# Multimodal Prediction of Breast Cancer Recurrence Assays and Risk of Recurrence

**DOI:** 10.1101/2022.07.07.499039

**Authors:** Frederick M. Howard, James Dolezal, Sara Kochanny, Galina Khramtsova, Jasmine Vickery, Andrew Srisuwananukorn, Anna Woodard, Nan Chen, Rita Nanda, Charles M. Perou, Olufunmilayo I. Olopade, Dezheng Huo, Alexander T. Pearson

## Abstract

Gene expression-based recurrence assays are strongly recommended to guide the use of chemotherapy in hormone receptor-positive, HER2-negative breast cancer, but such testing is expensive, can contribute to delays in care, and may not be available in low-resource settings. Here, we describe the training and independent validation of a deep learning model that predicts recurrence assay result and risk of recurrence using both digital histology and clinical risk factors. We demonstrate that this approach outperforms an established clinical nomogram (area under the receiver operating characteristic curve of 0.833 versus 0.765 in an external validation cohort, p = 0.003), and can identify a subset of patients with excellent prognoses who may not need further genomic testing.

## Main

Breast cancer is the leading cause of cancer death for women globally with an estimated 1.7 million cases diagnosed each year^1^. There is an unmet global clinical need for accurate diagnosis and treatment of breast cancer in response to the rising global burden of disease. Breast cancer is a biologically heterogeneous disease and genomic biomarkers have been developed to tailor therapeutic decisions. Hormone receptor-positive (HR+) breast cancer constitutes about 70% of newly diagnosed cases in the United States^2^, although lower rates are seen outside of western / European populations^3^. Gene expression-based recurrence score assays, such as OncotypeDx (ODX), MammaPrint (MP), Prosigna, and EndoPredict have been transformative for breast cancer management and are strongly recommended by National Comprehensive Cancer Network^4^ and American Society of Clinical Oncology (ASCO)^5^ guidelines to aid decisions regarding the use of chemotherapy. However, genomic testing is costly^6^, is underutilized in minorities and low resource settings^7^, and can take weeks to perform leading to significant delays in care^8^. Clinical nomograms have been developed to identify patients at high risk of recurrence, but do not obviate the need for genomic testing^9^. Compared to gene expression assays, hematoxylin and eosin (H&E) stained pathology images are readily available for all patients with cancer worldwide. Deep learning (DL) is a recent advance in the field of artificial intelligence (AI) which excels at quantitative image analysis. From histology, DL models can automatically identify high-level image features, which in turn can be used to predict outcomes of interest, such as tumor grade, gene expression, and genetic alterations^10–12^. As DL has been shown to predict breast cancer gene expression from tumor histology (including molecular subtype as well as genes involved in cell-cycle, angiogenesis, and immune response pathways)^10,11,13^, we hypothesized that DL can predict gene expression-based recurrence score assays directly from H&E stained pathology. A DL risk predictor which utilizes digital histology rather than requiring gene expression testing would have the potential to facilitate expedient decisions about prognosis and need for chemotherapy while reducing the cost of cancer care.

To develop an accurate DL model for prediction of recurrence score directly from histology, we used a framework of two consecutive modules applied to image tiles extracted from the digital slide – one to predict tumor likelihood and a second to predict likelihood of recurrence (Figure 1). The first DL module identified tumor regions of interest versus surrounding normal tissue using pathologist tumor annotations from n = 1,039 patients in The Cancer Genome Atlas (TCGA, Supplementary Table 1), achieving an average tile-level area under the receiver operating characteristic curve (AUROC) of 0.85 when assessed using internal three-fold cross-validation in TCGA. The second module was trained on image tiles from within the pathologist-annotated malignant areas from TCGA (n = 1,039 patients) to predict the results of recurrence assays (calculated using gene expression data). A DL pathology recurrence score prediction was obtained by weighting the tile-level recurrence score by tile-level tumor likelihood across all tiles to provide a patient-level prediction. Furthermore, to assess if integrating clinical data improves the discriminatory capacity of our model, we developed a combined model incorporating the DL pathologic prediction and a clinical predictor of high ODX scores. A logistic regression was fit within TCGA-BRCA using our DL model prediction and ODX prediction from a previously published clinical nomogram developed by researchers from the University of Tennessee^9^. This clinical nomogram incorporates patient age, tumor size, progesterone receptor (PR) status, tumor grade, and histologic subtype. In the HR+/HER2-subset of our training cohort from TCGA (n = 522, reflective of the population where ODX is usually performed), average AUROC for prediction of high ODX score was 0.776 (95% CI 0.595 – 0.913) for the DL pathology model, 0.768 (95% CI 0.648 – 0.868) for the Tennessee nomogram, and 0.816 (95% CI 0.688 – 0.913) for the combined model (Figure 2a).

**Figure 1.**
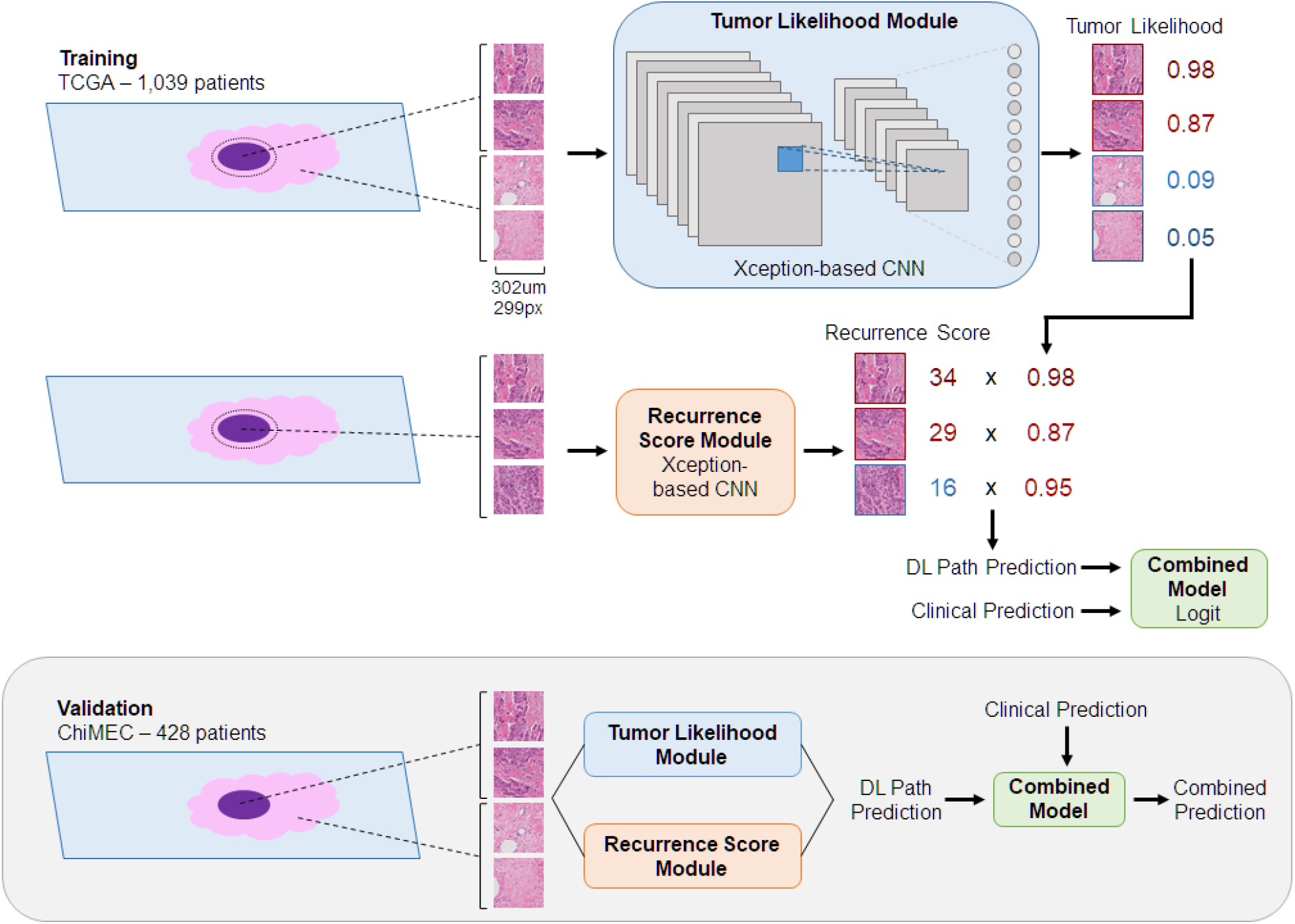
Overview of Study Design and Model Architecture. **a**. Two Xception-based deep learning models were trained on 1,039 patients from TCGA to allow for unsupervised predictions on external data. One model was trained to identify image tiles within pathologist annotation of tumor versus background image tiles. The second model was trained to predict a research version of the 21-gene recurrence score calculated from gene expression data from the annotated tumor regions from TCGA, and Bayesian optimization was used to select optimal model hyperparameters for recurrence score prediction. Finally, a combined clinical / pathologic model was developed by fitting a logistic regression to deep learning model predictions and the University of Tennessee clinical risk model. Performance of the deep learning pathologic, clinical, and combined models were then assessed in the external ChiMEC cohort (n = 428 for OncotypeDx, n = 88 for MammaPrint).

**Figure 2.**
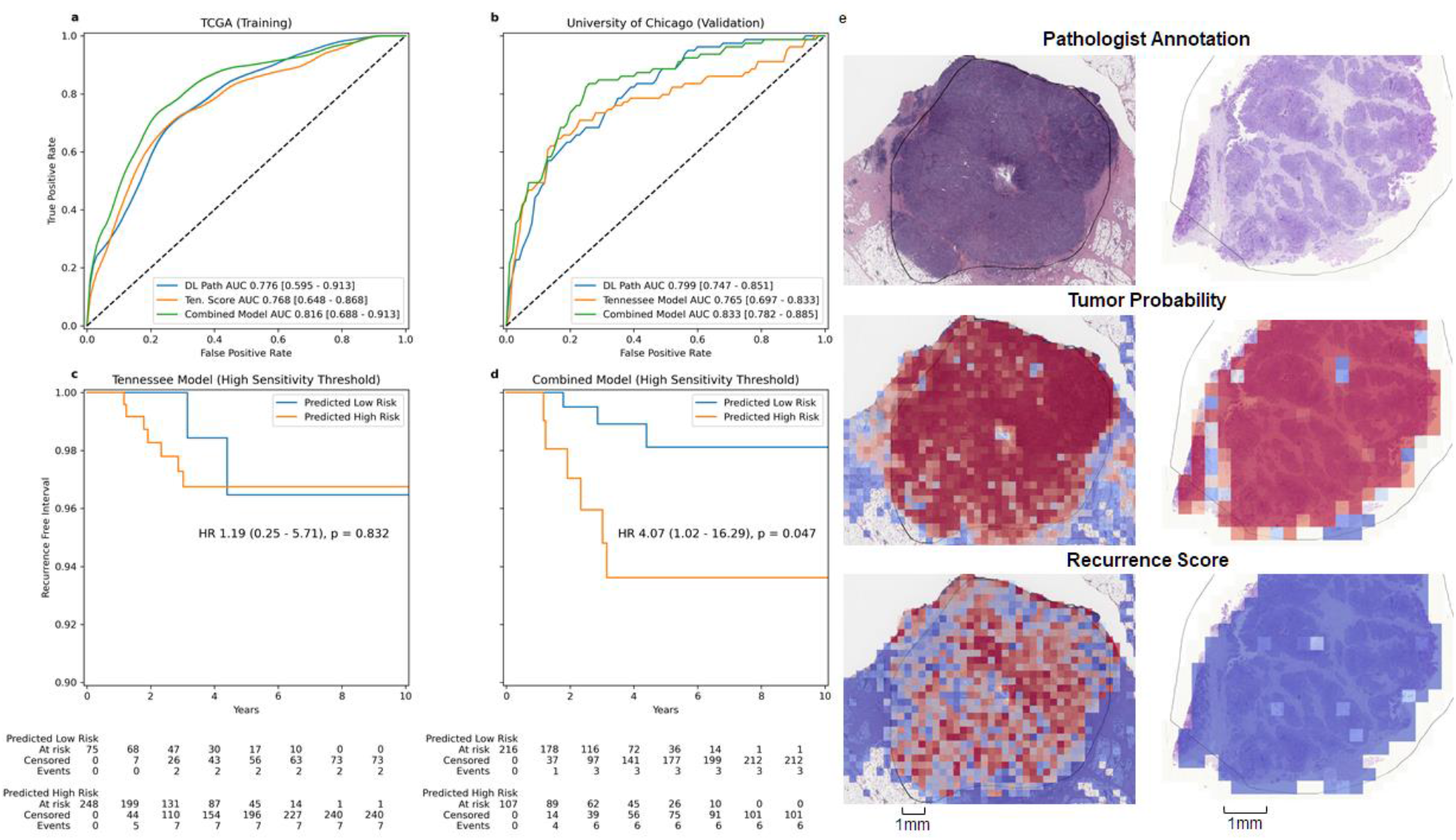
Model Results and Interpretation. **a**. Average patient-level AUROC for prediction of high-risk recurrence score in HR+/HER2-patients from TCGA (n = 522). **b**. Patient-level AUROC for prediction of high-risk recurrence score in the University of Chicago cohort (n = 455). **c & d**. Kaplan-Meier curves illustrate recurrence free interval with endocrine therapy alone in patients predicted to have a high-risk Oncotype score using high-sensitivity thresholds from the Tennessee Nomogram and combined predictive model identified from TCGA. **e**. On the left, heatmaps illustrate model predictions of tumor probability (middle) compared to pathologist annotation (top), and illustrate regions suggestive of high recurrence score (bottom) in a patient with a high recurrence score. To the right, similar heatmaps illustrate model predictions in a patient with a low recurrence score.

To validate these findings in an external cohort, we assessed model performance from the frozen DL pathologic and combined models trained on TCGA in n = 428 (Supplementary Table 2) patients from the University of Chicago Medical Center (UCMC) which had ODX testing performed and pathologic samples available. AUROC for prediction of high ODX score of the combined model was 0.833 (95% CI 0.782 – 0.885), which was significantly higher than either the DL pathology model (AUROC 0.799, 95% CI 0.747 – 0.851, p = 0.026) or the Tennessee nomogram (AUROC 0.765, 95% CI 0.697 – 0.833, p = 0.003, Figure 2b, Supplementary Table 3). The correlation coefficient between model predictions and numeric ODX score was also highest in the combined model (Pearson r = 0.545, 95% CI 0.475 – 0.608, Supplementary Figure 1). Performance was similar in Black and White patient subgroups (other racial/ethnic groups not assessed due to small sample size), with the combined model outperforming the clinical model in both subgroups (Supplementary Table 4).

As ODX was initially developed to predict prognosis in patients treated with endocrine therapy, we evaluated the prognostic accuracy of models in this subset of patients (n = 323) at UCMC. We modeled the prognostic accuracy using Cox proportional hazards models incorporating predictions from each model as a single variable, and compared the prognostic accuracy of each of these Cox models. Each model was significantly associated with recurrence-free interval (RFI, Supplementary Table 5), but the Harrell’s concordance index (C-index)^14^ was highest for the combined model (HR 2.02 per standard deviation, 95% CI 1.16 – 3.52, p = 0.013, C-index 0.751), nearly reaching the C-index of the actual ODX score (0.776). No model was associated with RFI among patients receiving chemotherapy when included as a single variable in a Cox model, which may be due to confounding variables influencing treatment decisions and the use of ODX to select patients for treatment. Finally, we compared the ability of the three models to perform as highly-sensitive rule-out tests, to identify patients who do not require ODX testing. We selected a threshold for each model that achieved a sensitivity of 95% in the TCGA HR+/HER2-cohort (Supplementary Table 6), and then applied that threshold to UCMC patients. The true sensitivities were lower in the UCMC cohort, but the specificity was higher for the combined model (sensitivity 84.8%, specificity 68.2%) than for the Tennessee nomogram (sensitivity 88.6%, specificity 21.8%) despite similar sensitivities. Additionally, recurrence-free interval among endocrine treated patients was higher in patients predicted to have low-risk ODX score with the UCMC model using the high sensitivity cutoff; which was not seen when using the Tennessee nomogram (Figure 2c-d).

We performed a similar analysis to evaluate DL as a predictor of high-risk MP scores. As there is not a widely used nomogram for high-risk MP prediction, we developed a clinical predictor from the National Cancer Data Base (NCDB). A combined model had numerically higher accuracy in the prediction of high-risk MP scores (AUROC 0.768, 95% CI 0.668 – 0.867) than a clinical model (AUROC 0.754, 95% CI 0.650 – 0.857, p = 0.662) or a pathologic model (AUROC 0.744, 95% CI 0.639 – 0.849, p = 0.517) in a validation cohort of n = 88 UCMC patients, but this did not reach statistical significance, perhaps in part due to the small sample size (Supplementary Figure 2, Supplementary Table 2). There was only one recurrence in the MP subgroup at UCMC, so prognostic comparisons to actual MP scores were not performed.

Finally, to help understand the nature of predictions made by this DL model, study pathologists independently reviewed heatmaps of the recurrence score module from 20 slides each with high-risk and low-risk predictions in the UCMC cohort. Notable features identified by heatmaps included necrosis (both comedonecrosis and coagulative necrosis), lymphovascular invasion, high-grade densely packed tumor nests, and infiltrative borders (Supplementary Figure 3). To further demonstrate the correlation of these features with model predictions, we compared predictions in out-of-sample cases in the TCGA cohort with and without selecting previously annotated histologic features. We found that pathologic prediction of high-risk ODX was associated with higher grade (p = 3.06 × 10^−85^), lymphovascular invasion (p = 0.023), and necrosis (p = 1.39 × 10^−62^, Supplementary Table 7).

The only other published DL method predicting recurrence scores from pathology used automated tubule nuclei quantification to differentiate high-grade tumors with ODX scores of > 30 versus low-grade tumors with ODX scores of < 18, demonstrating an AUROC of 0.76.^15^ However, this AUROC is likely artificially inflated as intermediate ODX scores of 18 - 30 were excluded, and this classifier has not been proven for categorization of high-grade / low ODX or low-grade / high ODX tumors. DL models have been deployed on small-scale datasets using only clinical and immunohistochemical features scored by pathologists^16,17^, but as we have described, DL on pathology can identify novel features that are independent from clinicopathologic features that are routinely available and the combined model developed here does not require special immunohistochemistry stains aside from PR status result. Strengths of this study include the consistency of performance in both training and validation subsets for model accuracy, as well as in racial/ethnic subgroups (which is essential given potential inequities in DL^18^), and result in meaningful improvements in accuracy and prognostic ability over existing clinical models. Additionally, the correlation of predictions with known high-risk histologic factors including grade, necrosis, and lymphovascular invasion suggest that biologically relevant features are identified by this weakly supervised DL pathologic approach. The models developed here are limited by the fact that patients from TCGA do not have clinical-grade recurrence assay results available, which may contribute to the decrease in sensitivity when attempting to identify a high-sensitivity threshold. However, model performance for ODX prediction was similar with both out-of-sample cross-validation in TCGA and in the external UCMC cohort, leading to confidence in this methodology. No model was prognostic in patients receiving chemotherapy, which may be due to confounding effects, but the small sample size limits multivariable models to control for clinical risk factors. Additionally, the prognostic value of all models was reduced when examining recurrence-free survival as opposed to recurrence-free interval, and larger sample sizes may be needed to confirm the prognostic value of our proposed model. Our tumor likelihood module slightly underperformed compared to other reports of breast cancer classification using DL^19,20^, but accuracy was influenced by pathologist annotations which sometimes did not precisely outline the tumor border (as seen in Figure 2e).

Understanding of the genomic features underlying cancer recurrence and chemotherapy benefit has evolved and is now a routine part of breast cancer care. ASCO recently added the development and integration of deep learning technology into cancer research as a priority in 2021^21^, as artificial intelligence has the potential to rectify disparities and supplement or improve genomic testing. This study illustrates the development of an effective DL biomarker that improves on existing clinical predictors of low recurrence risk tumors. ODX testing is estimated to grow in cost to $231 million annually in the USA^6^, and using a highly sensitive cutoff as described above could be used to limit testing in patients who are very unlikely to have positive results. Furthermore, given the heterogeneity of breast tumors, this methodology could be applied to multiple pathologic samples in a single patient to potentially increase confidence in results. With training on larger datasets with clinical-grade recurrence assays available to optimally tune thresholds, this approach could improve the speed at which treatment decisions are made due to the time-consuming nature of genomic testing, reduce the cost of care, and be utilized worldwide where genomic assays are not available.

## Methods

### Ethics Statement

All experiments were conducted in accordance with the Declaration of Helsinki and the study was approved by the University of Chicago Institutional Review Board, IRB 22-0707. For model training, patients were included from the TCGA breast cancer cohort (BRCA)^22^. For validation, anonymized archival tissue samples were retrieved from the University of Chicago from January 1^st^ 2006 through December 21^st^ 2020 where recurrence score results were available.

### Model Development

First, an automated tumor detection module was trained to distinguish breast tumor from background tissue in digitally scanned H&E slides. From TCGA, 1,099 slides were annotated manually by study pathologists to distinguish tumor from surrounding stroma. Tessellated image tiles were extracted from within areas of tumor with an edge length of 302 microns and downscaled to a width of 299 pixels, consistent with an optical resolution of 10x. Tile extraction and DL model training was performed with the Slideflow pipeline^23^, using an Xception^24^ convolutional neural network backbone with a variable number of fully connected hidden layers prior to outcome prediction. The tumor likelihood module was trained with hyperparameters as listed in Supplementary Table 8 to distinguish tiles originating from within the tumor annotation from those outside the annotation. Model performance was assessed with average accuracy over three cross-fold validation, and a separate model was trained on the entire dataset for prediction on external patients.

Next, a separate DL module was trained to predict recurrence score from tumor image tiles. As the clinically validated multigene recurrence assay results are not available from TCGA, “research-based” versions of ODX and MP were calculated using the statistical methods from the published development of these assays^25–27^ applied to mRNA expression data from TCGA. To determine a threshold for high-risk “research-based” ODX score, the 15^th^ percentile result of HR+/HER2-patients in TCGA was used, as this is the percentile of patients with ODX score of 26 or higher in the National Cancer Database^9^. In the University of Chicago cohort, we used standard high-risk cutpoints of ODX score of 26 or higher, and MP score of lower than 0. Hyperparameters for these models were chosen with Bayesian optimization of cross validated tile level AUROC, run over 50 iterations (Supplemental Table 8, Supplementary Figure 4). Two sets of three cross folds were used for optimization, and folds were generated with site preservation^28^ to maximize generalizability. Patient-level predictions were calculated by weighting the average of tile-level predictions from this recurrence score prediction module according to a tile’s likelihood of tumor from the first module.

For clinical prediction of recurrence the University of Tennessee Nomogram^9^ was computed for each patient in TCGA; grade is not available in the original TCGA annotations but has been assessed and reported in prior work^29^. Precise tumor size was not provided in TCGA but was estimated from tumor stage group. Mean imputation was used when input data for the nomogram was not available to maximize included cases for the development of a combined model. Finally, a logistic regression was fit using the out-of-sample prediction from the pathologic model combined with the prediction from the clinical nomogram. Thresholds for computing model sensitivity were determined from TCGA (using interpolation to achieve an exact estimated sensitivity of 95%) and applied to the validation dataset from the University of Chicago.

Development of the MP prediction model proceeded in a similar fashion with a few key differences. As no widely used clinical model was available, we developed a clinical predictor from n = 6,938 non-metastatic HR+/HER2-patients from NCDB who were diagnosed with breast cancer between 2010 and 2017 and had MP testing results available. We used sequential forward feature selection to identify features that improved the AUROC for MP prediction in a logistic regression with 10-fold cross-validation, ultimately identifying grade, tumor size, PR status, lymphovascular invasion, ductal, mucinous, metaplastic, or medullary histology, and Black or Asian race for inclusion. A logistic regression incorporating these features was fit on all available data and used for prediction. We used the same optimized hyperparameters from ODX prediction for our DL pathologic MP model.

### Statistical Analysis

Internal validation of model accuracy for recurrence score prediction in TCGA was estimated by averaging patient-level AUROC over three-fold site preserved cross-validation, and 1000x bootstrapping for confidence interval estimation. External validation was performed with single fixed models generated from all TCGA data, using Delong’s method for estimation of AUROC confidence intervals and comparison^30^. The prognostic accuracy of models for RFI was assessed with the Wald test in univariable Cox models. Two sided t-tests were performed to compare DL pathologic model predictions between patients with or without select pathologic features. All statistical analysis was performed in Python 3.8, Lifelines 0.27.0, and Scipy 1.8.0 and performed at the α = 0.05 significance level. Given the limited number of statistical tests, performed in different subsets of patients, and the exploratory nature of this work, correction for multiple hypothesis testing was not performed.

## Data Availability

Data from TCGA including digital histology and the clinical and genetic annotations used are available from https://portal.gdc.cancer.gov/ and https://cbioportal.org. The NCDB PUF is a HIPAA-compliant data file, which is made available to investigators from CoC-accredited cancer programs who complete an application process. Trained models evaluated in this paper and anonymized patient level annotations can be obtained at doi.org/10.5281/zenodo.6792391. All other data used in this study is available from authors upon reasonable request.

## Code Availability

Code utilized in model development and assessment is available at github.com/fmhoward/DLRS.

## Acknowledgments

F.M.H. received support from ASCO/CCF (2022YIA-6675470300) and the NIH/NCI (F32CA265232). A.T.P received support from the NIH/NIDCR (K08-DE026500), the NCI (U01-CA243075), the Adenoid Cystic Carcinoma Research Foundation, the Cancer Research Foundation, and the American Cancer Society. A.T.P. and D.H. received support from the Department of Defense (BC211095P1). D.H., R.N., and O.I.O. received support from the NIH/NCI (1P20-CA233307). O.I.O. received support from Susan G Komen (SAC 210203). D.H. and O.I.O received support from Breast Cancer Research Foundation (BCRF-21-071). C.M.P. was supported by funds from the NCI Breast SPORE program (P50-CA058223), and the Breast Cancer Research Foundation. We would like to thank Shirley A. Mertz of the Metastatic Breast Cancer Alliance for feedback on this study.

## Author Contributions

F.M.H. and A.T.P. were responsible for concept proposal and study design. F.M.H., J.D., S.K., and A.S. performed essential programming work. G.K. and J.V. performed manual oversight and quality control for digital pathology, along with segmentation of tumor. F.M.H., J.D., S.K., A.S., A.W., O.I.O., D.H. and A.T.P. contributed to data interpretation and statistical approaches. All authors contributed to the data analysis and writing of the manuscript.

## References

1. Bray F, Ferlay J, Soerjomataram I, Siegel RL, Torre LA, Jemal A. Global cancer statistics 2018: GLOBOCAN estimates of incidence and mortality worldwide for 36 cancers in 185 countries. CA: A Cancer Journal for Clinicians. 2018;68(6):394–424. doi:10.3322/caac.21492

2. Brenton JD, Carey LA, Ahmed AA, Caldas C. Molecular classification and molecular forecasting of breast cancer: ready for clinical application? J Clin Oncol. 2005;23(29):7350–7360. doi:10.1200/JCO.2005.03.3845

3. Huo D, Ikpatt F, Khramtsov A, et al. Population Differences in Breast Cancer: Survey in Indigenous African Women Reveals Over-Representation of Triple-Negative Breast Cancer. J Clin Oncol. 2009;27(27):4515–4521. doi:10.1200/JCO.2008.19.6873

4. Gradishar WJ, Anderson BO, Abraham J, et al. Breast Cancer, Version 3.2020, NCCN Clinical Practice Guidelines in Oncology. J Natl Compr Canc Netw. 2020;18(4):452–478. doi:10.6004/jnccn.2020.0016

5. Andre F, Ismaila N, Henry NL, et al. Use of Biomarkers to Guide Decisions on Adjuvant Systemic Therapy for Women With Early-Stage Invasive Breast Cancer: ASCO Clinical Practice Guideline Update-Integration of Results From TAILORx. J Clin Oncol. 2019;37(22):1956–1964. doi:10.1200/JCO.19.00945

6. Mariotto A, Jayasekerea J, Petkov V, et al. Expected Monetary Impact of Oncotype DX Score-Concordant Systemic Breast Cancer Therapy Based on the TAILORx Trial. J Natl Cancer Inst. 2019;112(2):154–160. doi:10.1093/jnci/djz068

7. Press DJ, Ibraheem A, Dolan ME, Goss KH, Conzen S, Huo D. Racial disparities in omission of oncotype DX but no racial disparities in chemotherapy receipt following completed oncotype DX test results. Breast Cancer Res Treat. 2018;168(1):207–220. doi:10.1007/s10549-017-4587-8

8. Losk K, Vaz-Luis I, Camuso K, et al. Factors Associated With Delays in Chemotherapy Initiation Among Patients With Breast Cancer at a Comprehensive Cancer Center. J Natl Compr Canc Netw. 2016;14(12):1519–1526. doi:10.6004/jnccn.2016.0163

9. Orucevic A, Bell JL, King M, McNabb AP, Heidel RE. Nomogram update based on TAILORx clinical trial results - Oncotype DX breast cancer recurrence score can be predicted using clinicopathologic data. The Breast. 2019;46:116–125. doi:10.1016/j.breast.2019.05.006

10. Kather JN, Heij LR, Grabsch HI, et al. Pan-cancer image-based detection of clinically actionable genetic alterations. Nature Cancer. Published online July 27, 2020:1-11. doi:10.1038/s43018-020-0087-6

11. Couture HD, Williams LA, Geradts J, et al. Image analysis with deep learning to predict breast cancer grade, ER status, histologic subtype, and intrinsic subtype. npj Breast Cancer. 2018;4(1):1–8. doi:10.1038/s41523-018-0079-1

12. Campanella G, Hanna MG, Geneslaw L, et al. Clinical-grade computational pathology using weakly supervised deep learning on whole slide images. Nature Medicine. 2019;25(8):1301–1309. doi:10.1038/s41591-019-0508-1

13. Schmauch B, Romagnoni A, Pronier E, et al. A deep learning model to predict RNA-Seq expression of tumours from whole slide images. Nature Communications. 2020;11(1):3877. doi:10.1038/s41467-020-17678-4

14. Liao JJZ, Lewis JW. A Note on Concordance Correlation Coefficient. PDA Journal of Pharmaceutical Science and Technology. 2000;54(1):23–26.

15. Romo-Bucheli D, Janowczyk A, Gilmore H, Romero E, Madabhushi A. Automated Tubule Nuclei Quantification and Correlation with Oncotype DX risk categories in ER+ Breast Cancer Whole Slide Images. Sci Rep. 2016;6(1):32706. doi:10.1038/srep32706

16. Baltres A, Al Masry Z, Zemouri R, et al. Prediction of Oncotype DX recurrence score using deep multi-layer perceptrons in estrogen receptor-positive, HER2-negative breast cancer. Breast Cancer. 2020;27(5):1007–1016. doi:10.1007/s12282-020-01100-4

17. Kim I, Choi HJ, Ryu JM, et al. A predictive model for high/low risk group according to oncotype DX recurrence score using machine learning. Eur J Surg Oncol. 2019;45(2):134–140. doi:10.1016/j.ejso.2018.09.011

18. Char DS, Shah NH, Magnus D. Implementing Machine Learning in Health Care — Addressing Ethical Challenges. N Engl J Med. 2018;378(11):981–983. doi:10.1056/NEJMp1714229

19. Aresta G, Araújo T, Kwok S, et al. BACH: Grand challenge on breast cancer histology images. Med Image Anal. 2019;56:122–139. doi:10.1016/j.media.2019.05.010

20. Yan R, Ren F, Wang Z, et al. Breast cancer histopathological image classification using a hybrid deep neural network. Methods. 2020;173:52–60. doi:10.1016/j.ymeth.2019.06.014

21. Smith SM, Wachter K, Burris HA, et al. Clinical Cancer Advances 2021: ASCO’s Report on Progress Against Cancer. JCO. 2021;39(10):1165–1184. doi:10.1200/JCO.20.03420

22. Comprehensive molecular portraits of human breast tumors. Nature. 2012;490(7418):61–70. doi:10.1038/nature11412

23. Dolezal J, Kochanny S, Howard F. Jamesdolezal/Slideflow: Slideflow 1.0 - Official Public Release. Zenodo; 2021. doi:10.5281/zenodo.5708490

24. Chollet F. Xception: Deep Learning with Depthwise Separable Convolutions. 161002357 [cs]. Published online April 4, 2017. Accessed August 12, 2020. http://arxiv.org/abs/1610.02357

25. Paik S, Shak S, Tang G, et al. A multigene assay to predict recurrence of tamoxifen-treated, node-negative breast cancer. N Engl J Med. 2004;351(27):2817–2826. doi:10.1056/NEJMoa041588

26. van ‘t Veer LJ, Dai H, van de Vijver MJ, et al. Gene expression profiling predicts clinical outcome of breast cancer. Nature. 2002;415(6871):530–536. doi:10.1038/415530a

27. van de Vijver MJ, He YD, van ‘t Veer LJ, et al. A Gene-Expression Signature as a Predictor of Survival in Breast Cancer. New England Journal of Medicine. 2002;347(25):1999–2009. doi:10.1056/NEJMoa021967

28. Howard FM, Dolezal J, Kochanny S, et al. The impact of site-specific digital histology signatures on deep learning model accuracy and bias. Nat Commun. 2021;12(1):1–13. doi:10.1038/s41467-021-24698-1

29. Thennavan A, Beca F, Xia Y, et al. Molecular analysis of TCGA breast cancer histologic types. Cell Genom. 2021;1(3):100067. doi:10.1016/j.xgen.2021.100067

30. DeLong ER, DeLong DM, Clarke-Pearson DL. Comparing the areas under two or more correlated receiver operating characteristic curves: a nonparametric approach. Biometrics. 1988;44(3):837–845.

