## Supplementary Materials for "Multimodal Prediction of Breast Cancer Recurrence Assays and Risk of Recurrence"

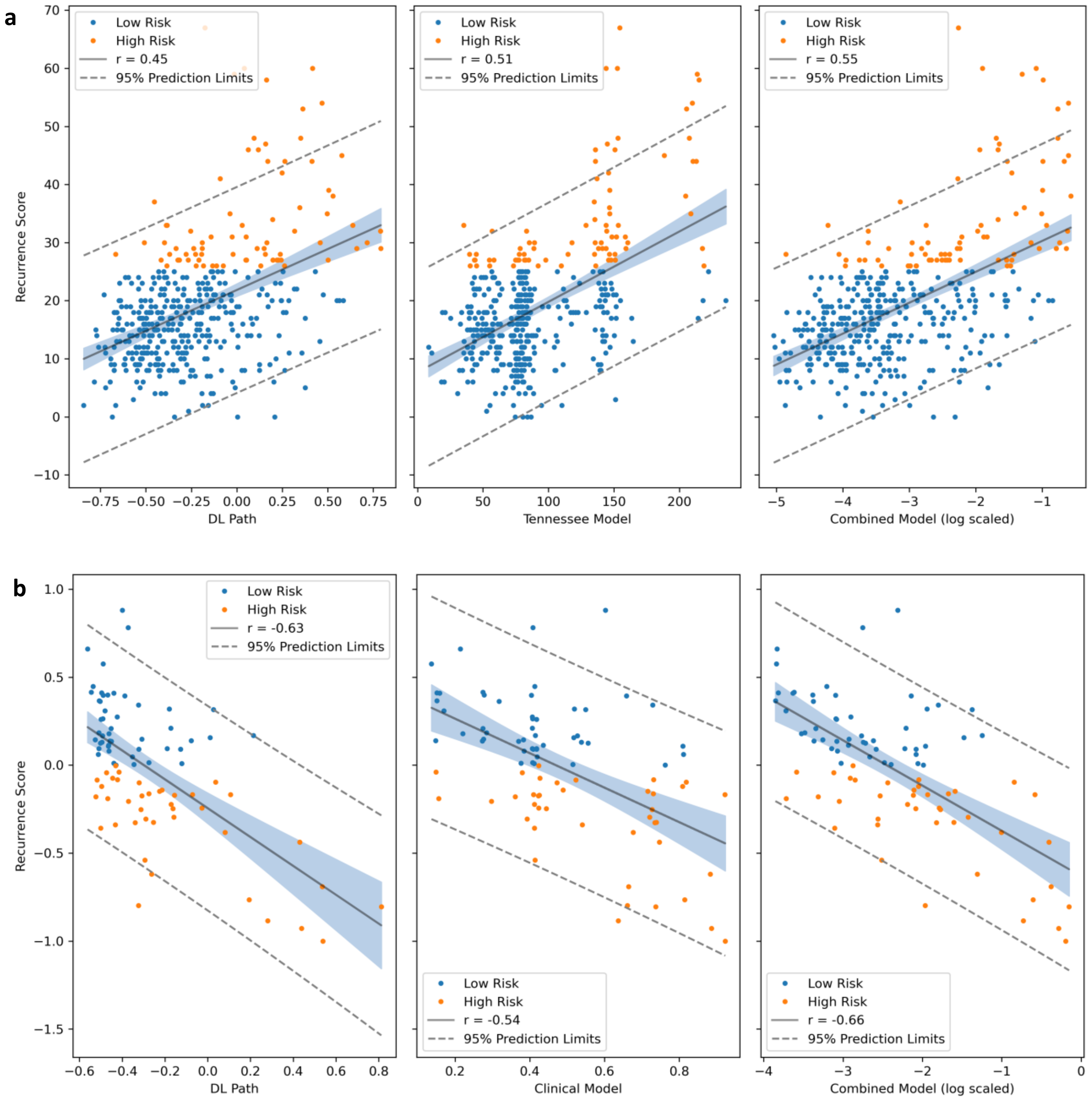


**Supplementary Figure 1. Linear Correlation of Models Predictions with OncotypeDx and MammaPrint Scores. a**. Correlation of the deep learning pathology, Tennessee, and combined models with OncotypeDx scores in the University of Chicago Cohort (n = 428). **b**. Correlation of the deep learning pathology, clinical, and combined models with MammaPrint scores in the University of Chicago Cohort (n = 88). **Abbreviations: DL = Deep Learning.**

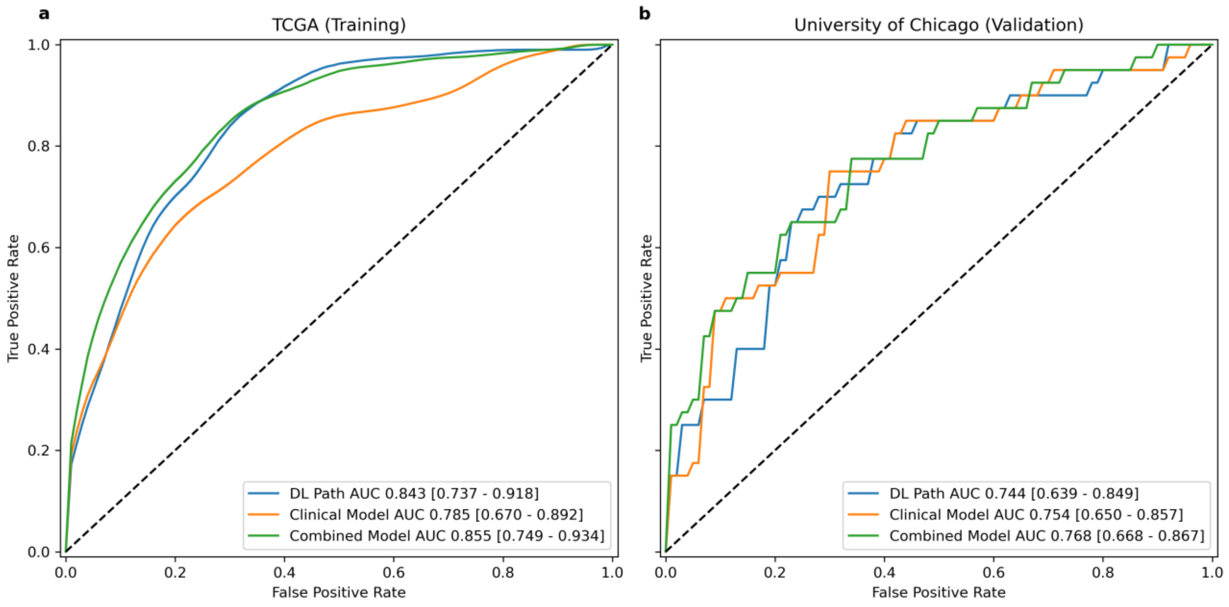


**Supplementary Figure 2. Predictive Accuracy of MammaPrint Prediction Models. a.** Receiver operating characteristic curves for MammaPrint prediction for the deep learning pathologic, clinical, and combined models in HR/HER2- patients from TCGA (n = 522). **b.** The same curves plotted for the external University of Chicago cohort (n = 88). **Abbreviations: DL = Deep Learning. AUC = Area Under the Receiver Operating Characteristic Curve.**

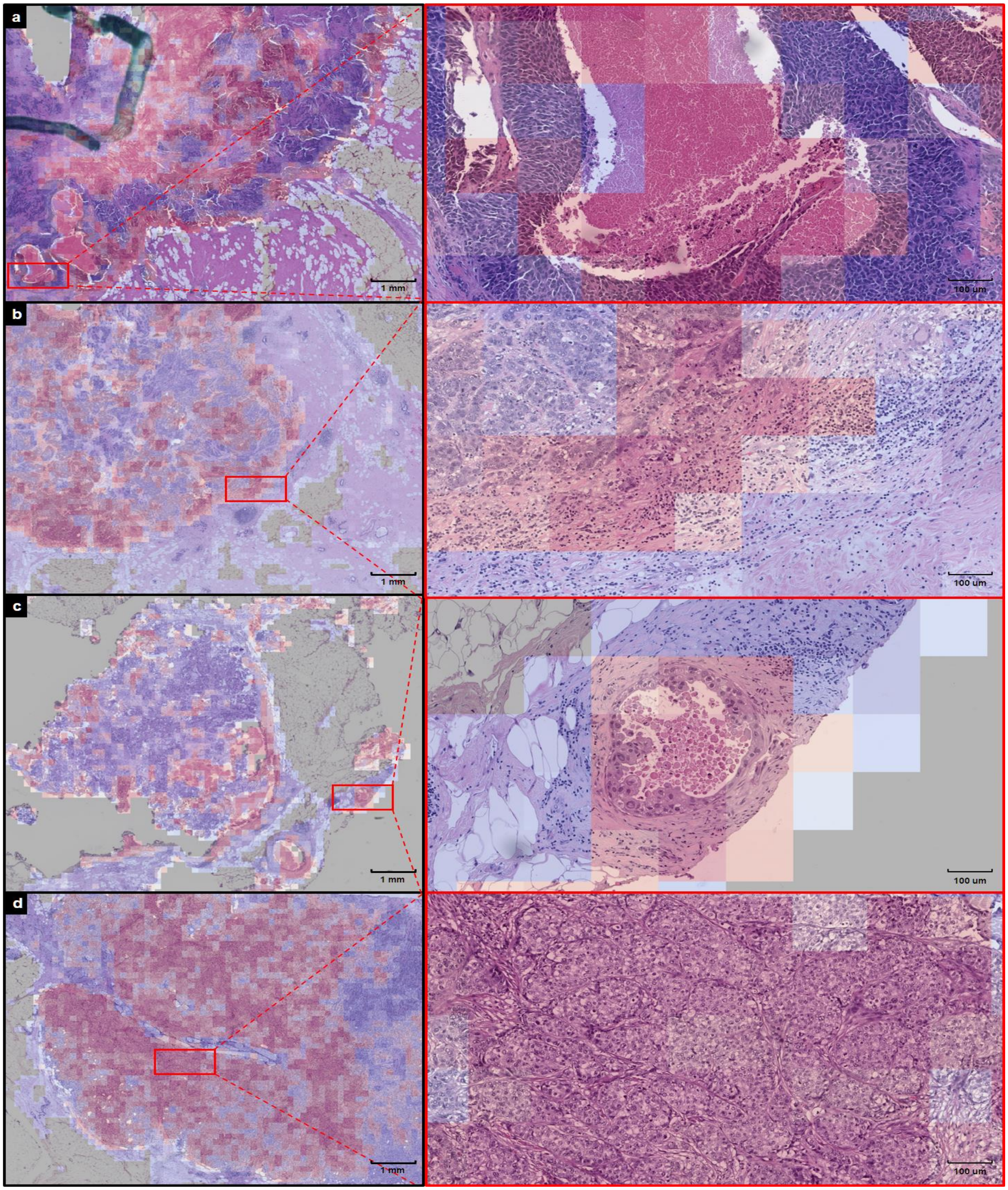


**Supplementary Figure 3. Heatmaps of the Recurrence Score Prediction Model on Select High-Risk Tumors.** Review of patients with predicted high recurrence scores identified several pathologic characteristics of tumors predicted to be high risk of recurrence including **a.** Comedo necrosis, **b.** infiltrative borders, **c.** lymphovascular invasion, and **d.** high grade densely packed tumor nests.

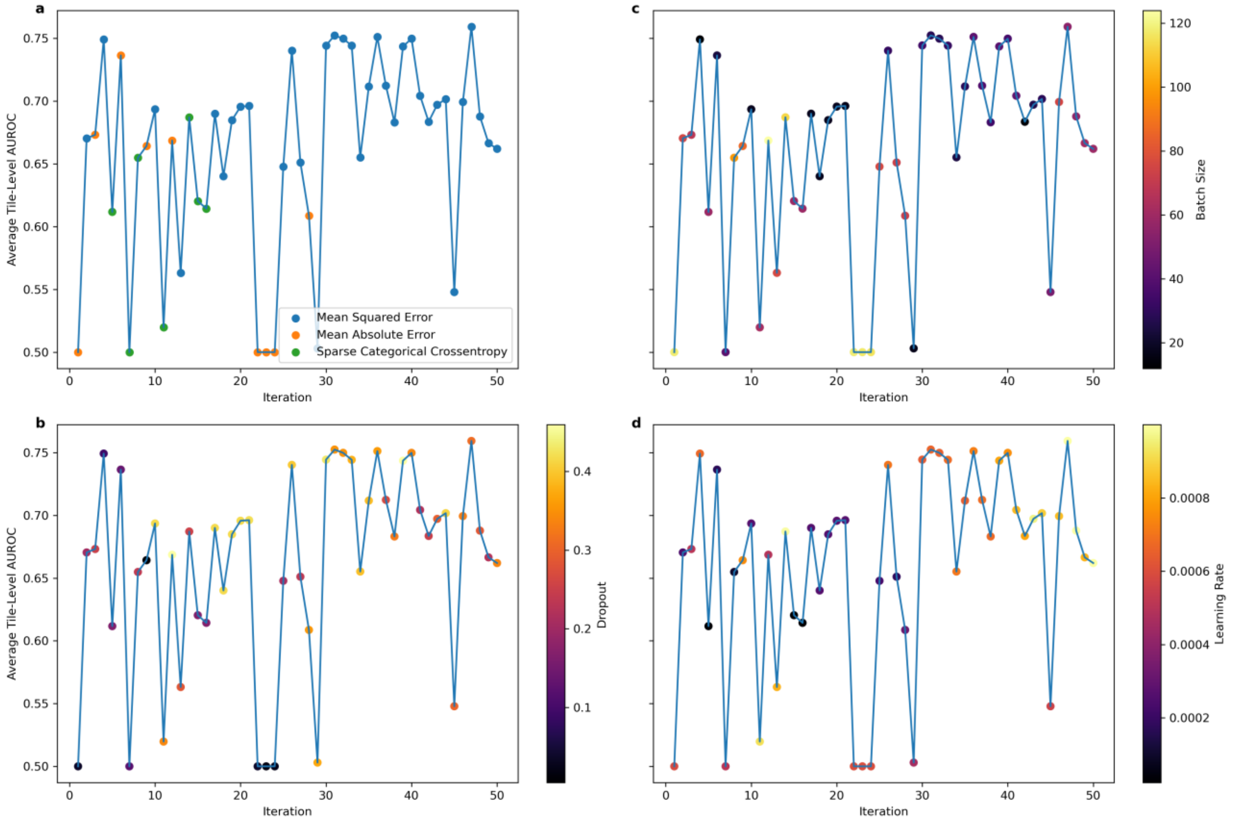


**Supplementary Figure 4. Tile-Level AUROC for Prediction of High-Risk OncotypeDx Score During Hyperparameter Optimization.** Results are listed over 50 iterations of Bayesian optimization with objective of maximizing average tile-level AUROC over two replicates of three site-preserved cross folds in The Cancer Genome Atlas. Plots highlight the associations of selected hyperparameters **a.** loss function, **b.** batch size, **c.** dropout, and **d.** learning rate with model performance. **Abbreviations: AUROC = Area Under the Receiver Operating Characteristic Curve.**

**Supplementary Table 1. Baseline Demographics from Cohorts from The Cancer Genome Atlas used for Model Training.**

| | | Missing | Overall |
|---|---|---|---|
| **n** | | | 1039 |
| **Age, mean (SD)** | | 0 | 58.6 (13.2) |
| **Sex, n (%)** | **Female** | 0 | 1027 (98.8) |
| | **Male** | | 12 (1.2) |
| **Race, n (%)** | **Asian** | 94 | 60 (6.3) |
| | **Black** | | 162 (17.1) |
| | **Other** | | 1 (0.1) |
| | **White** | | 722 (76.4) |
| **Ethnicity, n (%)** | **Hispanic** | 166 | 38 (4.4) |
| | **Non-Hispanic** | | 835 (95.6) |
| **Histologic Subtype, n (%)** | **Ductal** | 0 | 632 (60.8) |
| | **Ductal and Lobular** | | 114 (11.0) |
| | **Lobular** | | 176 (16.9) |
| | **Other** | | 117 (11.3) |
| **Grade, n (%)** | **1** | 2 | 229 (22.1) |
| | **2** | | 428 (41.3) |
| | **3** | | 380 (36.6) |
| **Tumor Size (mm), mean (SD)** | | 0 | 31.8 (12.1) |
| **Nodal Status, n (%)** | **Negative** | 19 | 488 (47.8) |
| | **Positive** | | 532 (52.2) |
| **ER Status, n (%)** | **Negative** | 49 | 224 (22.6) |
| | **Positive** | | 766 (77.4) |
| **PR Status, n (%)** | **Negative** | 52 | 325 (32.9) |
| | **Positive** | | 662 (67.1) |
| **Research-Only OncotypeDx Score, mean (SD)** | | 0 | 0.1 (1.0) |
| **Research-Only MammaPrint Score, mean (SD)** | | 0 | -0.0 (1.0) |
| **Chemotherapy, n (%)** | **No** | 1 | 919 (88.5) |
| | **Yes** | | 119 (11.5) |
| **Follow-up, mean (SD)** | | 1 | 3.5 (3.3) |
| **Recurrence, n (%)** | **Disease Free** | 0 | 902 (86.8) |
| | **Recurred** | | 137 (13.2) |
| **Vital Status, n (%)** | **Alive** | 0 | 893 (85.9) |
| | **Dead** | | 146 (14.1) |

**Supplementary Table 2. Baseline Demographics from Cohorts from the University of Chicago used for Model Validation.**

| | | Missing | OncotypeDx Cohort | Missing | MammaPrint Cohort |
|---|---|---|---|---|---|
| n | | - | 428 | - | 88 |
| Age, mean (SD) | | 0 | 56.3 (10.6) | 0 | 54.0 (12.7) |
| Sex, n (%) | Female | 0 | 424 (99.1) | 0 | 88 (100.0) |
| | Male | | 4 (0.9) | | 0 (0.0) |
| Race, n (%) | Asian | 8 | 27 (6.4) | 1 | 3 (3.4) |
| | Black | | 102 (24.3) | | 23 (26.4) |
| | Other | | 2 (0.5) | | 1 (1.1) |
| | White | | 289 (68.8) | | 60 (69.0) |
| Ethnicity, n (%) | Hispanic | 5 | 13 (3.1) | 2 | 5 (5.8) |
| | Non-Hispanic | | 410 (96.9) | | 81 (94.2) |
| Histologic Subtype, n (%) | Ductal | 0 | 303 (70.8) | 0 | 68 (77.3) |
| | Ductal and Lobular | | 40 (9.3) | | 9 (10.2) |
| | Lobular | | 71 (16.6) | | 10 (11.4) |
| | Other | | 14 (3.3) | | 1 (1.1) |
| Grade, n (%) | 1.0 | 0 | 69 (16.1) | 0 | 10 (11.4) |
| | 2.0 | | 279 (65.2) | | 58 (65.9) |
| | 3.0 | | 80 (18.7) | | 20 (22.7) |
| Tumor Size (mm), mean (SD) | | 0 | 21.0 (16.2) | 0 | 25.5 (23.1) |
| Nodal Status, n (%) | Negative | 0 | 363 (84.8) | 0 | 15 (17.0) |
| | Positive | | 65 (15.2) | | 73 (83.0) |
| ER Status, n (%) | Negative | 1 | 9 (2.1) | 0 | 7 (8.0) |
| | Positive | | 418 (97.9) | | 81 (92.0) |
| PR Status, n (%) | Negative | 0 | 53 (12.4) | 0 | 12 (13.6) |
| | Positive | | 375 (87.6) | | 76 (86.4) |
| HER2 Status, n (%) | Negative | 9 | 411 (98.1) | 4 | 83 (98.8) |
| | Positive | | 8 (1.9) | | 1 (1.2) |
| OncotypeDx Score, mean (SD) | | 0 | 18.6 (10.1) | 0 | Mammaprint Score, mean (SD) -0.0 (0.4) |
| Chemotherapy, n (%) | No | 0 | 324 (75.7) | 0 | 49 (55.7) |
| | Yes | | 104 (24.3) | | 39 (44.3) |
| Years Follow-up, mean (SD) | | 0 | 6.8 (4.0) | 0 | 3.2 (2.4) |
| Recurrence, n (%) | Disease Free | 2 | 411 (96.5) | 3 | 84 (98.8) |
| | Recurred | | 15 (3.5) | | 1 (1.2) |
| Vital Status, n (%) | Alive | 0 | 408 (95.3) | 0 | 86 (100.0) |
| | Dead | | 20 (4.7) | | 0 (0.0) |

**Supplementary Table 3. Predictive Accuracy for Recurrence Score.** Results are listed for correlation with numeric recurrence score, and AUROC for prediction of high-risk recurrence score. For the validation dataset, the AUROC of the combined model was compared to the individual component pathologic / clinical models using Delong's method. **Abbreviations: TCGA = The Cancer Genome Atlas. AUROC = Area Under the Receiver Operating Characteristic Curve)**

| | TCGA (Training) | | University of Chicago (Validation) | | | |
|---|---|---|---|---|---|---|
| | **Pearson Correlation Coefficient** | **AUROC** | **Pearson Correlation Coefficient** | **AUROC** | **z-statistic** | **p – value (AUROC compared to Combined Model)** |
| | | | **OncotypeDx** | | | |
| **Pathologic Model** | 0.506 (0.442 - 0.565) | 0.776 (0.595 - 0.913) | 0.448 (0.369 - 0.521) | 0.799 (0.747 - 0.851) | 2.227 | 0.026 |
| **Tennessee Clinical Nomogram** | 0.494 (0.429 - 0.554) | 0.768 (0.648 - 0.868) | 0.508 (0.435 - 0.576) | 0.765 (0.697 - 0.833) | 2.94 | 0.003 |
| **Combined Model** | 0.555 (0.495 - 0.610) | 0.816 (0.688 - 0.913) | 0.545 (0.475 - 0.608) | 0.833 (0.782 - 0.885) | - | - |
| | | | **MammaPrint** | | | |
| **Pathologic Model** | -0.622 (-0.670 - -0.568) | 0.843 (0.737 - 0.918) | -0.627 (-0.740 - -0.481) | 0.744 (0.639 - 0.849) | 0.648 | 0.517 |
| **Clinical Model (from NCDB)** | -0.525 (-0.583 - -0.462) | 0.785 (0.670 - 0.892) | -0.545 (-0.677 - -0.379) | 0.754 (0.650 - 0.857) | 0.437 | 0.662 |
| **Combined Model** | -0.648 (-0.694 - -0.597) | 0.855 (0.749 - 0.934) | -0.661 (-0.764 - -0.524) | 0.768 (0.668 - 0.867) | - | - |

**Supplementary Table 4. Predictive Accuracy in Racial Subgroups.** Results are listed for correlation with numeric recurrence score, and AUROC for prediction of high-risk recurrence score in the validation dataset for White (n = 308) and Black (n = 109), demonstrating that performance is preserved in racial subgroups.

**Abbreviations: AUROC = Area Under the Receiver Operating Characteristic Curve.**

| | **White Race** | | | **Black Race** | | |
|---|---|---|---|---|---|---|
| | **Pearson Correlation Coefficient** | **AUROC** | **p – value (AUROC compared to Combined Model)** | **Pearson Correlation Coefficient** | **AUROC** | **p – value (AUROC compared to Combined Model)** |
| **Pathologic Model** | 0.404 (0.303 - 0.496) | 0.779 (0.710 - 0.847) | 0.134 | 0.542 (0.388 - 0.666) | 0.828 (0.740 - 0.917) | 0.208 |
| **Tennessee Clinical Nomogram** | 0.488 (0.395 - 0.572) | 0.732 (0.636 - 0.827) | 0.013 | 0.549 (0.397 - 0.672) | 0.791 (0.691 - 0.892) | 0.063 |
| **Combined Model** | 0.512 (0.422 - 0.593) | 0.811 (0.740 - 0.883) | - | 0.628 (0.494 - 0.733) | 0.861 (0.783 - 0.940) | - |

**Supplementary Table 5. Prognostic Value of Models.** Results are listed for Cox proportional hazard models using the specified variable as the only input, for patients receiving endocrine therapy alone and for the whole dataset. Hazard ratios are computed using per standard deviation of input data given the different scales of the various models.

| | Patients Receiving Endocrine Therapy Alone (n = 323) | | | | Patients Receiving Chemotherapy (n = 103) | | | |
|---|---|---|---|---|---|---|---|---|
| | Hazard Ratio (95% CI) per unit SD | z-statistic | p-value | C-Index | Hazard Ratio (95% CI) per unit SD | z-statistic | p-value | C-Index |
| **Recurrence-Free Interval** | | | | | | | | |
| **Pathologic Model** | 1.724 (1.009 - 2.948) | 1.991 | 0.046 | 0.708 | 0.904 (0.38 - 2.151) | -0.229 | 0.819 | 0.5 |
| **Tennessee Clinical Nomogram** | 1.749 (1.087 - 2.814) | 2.306 | 0.021 | 0.68 | 1.204 (0.565 - 2.568) | 0.48 | 0.631 | 0.593 |
| **Combined Model (Log Scale)** | 2.023 (1.162 - 3.521) | 2.492 | 0.013 | 0.751 | 1.061 (0.47 - 2.396) | 0.143 | 0.886 | 0.495 |
| **OncotypeDx** | 1.853 (1.324 - 2.593) | 3.598 | 0.0003 | 0.776 | 1.429 (0.723 - 2.823) | 1.027 | 0.305 | 0.628 |
| **Recurrence-Free Survival** | | | | | | | | |
| **Pathologic Model** | 1.102 (0.736 - 1.649) | 0.47 | 0.638 | 0.562 | 0.825 (0.365 - 1.861) | -0.465 | 0.642 | 0.532 |
| **Tennessee Clinical Nomogram** | 1.13 (0.759 - 1.683) | 0.604 | 0.546 | 0.57 | 1.203 (0.595 - 2.432) | 0.514 | 0.607 | 0.595 |
| **Combined Model (Log Scale)** | 1.133 (0.754 - 1.704) | 0.602 | 0.547 | 0.574 | 1.006 (0.472 - 2.146) | 0.016 | 0.987 | 0.506 |
| **OncotypeDx** | 1.447 (1.052 - 1.99) | 2.272 | 0.023 | 0.648 | 1.399 (0.736 - 2.658) | 1.025 | 0.306 | 0.636 |

**Supplementary Table 6. Utility of Models as Rule-Out Tests.** Using TCGA data, thresholds were averaged over threefold cross validation to yield sensitivities of 95% for high-risk recurrence scores (with linear interpolation to achieve exact result of 95%). These thresholds were applied to the University of Chicago dataset to evaluate performance characteristics, and assess accuracy for prediction of recurrence (measured as hazard ratio for recurrence-free interval). **Abbreviations: TCGA = The Cancer Genome Atlas. Sen = Sensitivity. Spe = Specificity. PPV = Positive Predictive Value. NPV = Negative Predictive Value.**

| | TCGA | | | | University of Chicago | | | | | |
|---|---|---|---|---|---|---|---|---|---|---|
| | **Sen (%)** | **Spe (%)** | **PPV (%)** | **NPV (%)** | **Sen (%)** | **Spe (%)** | **PPV (%)** | **NPV (%)** | **Hazard Ratio (95% CI)** | **Hazard Ratio, No Chemo (95% CI)** |
| **Pathologic Model** | 95.0 | 29.0 | 17.3 | 97.3 | 82.3 | 61.3 | 32.5 | 93.9 | 2.041 (0.697 - 5.976) | 2.949 (0.737 - 11.799) |
| **Tennessee Clinical Nomogram** | 95.0 | 23.8 | 16.2 | 96.8 | 88.6 | 21.8 | 20.4 | 89.4 | 1.127 (0.318 - 3.996) | 1.186 (0.246 - 5.71) |
| **Combined Model** | 95.0 | 30.7 | 18.4 | 96.8 | 84.8 | 68.2 | 37.6 | 95.2 | 2.713 (0.927 - 7.938) | 4.074 (1.019 - 16.292) |

**Supplementary Table 7. Pathologic Characteristics of Predicted High Recurrence Score Patients.** In slides from The Cancer Genome Atlas, model predictions for high recurrence score were correlated with known pathologic features.

| Feature | n Slides | t-statistic | p-value |
|---|---|---|---|
| Necrosis | 1076 | 17.845 | $1.39 \times 10^{-62}$ |
| Lymphovascular Invasion | 1049 | 2.269 | 0.0234 |
| Grade 3 (versus grade 2 or 1) | 1096 | 21.414 | $3.06 \times 10^{-85}$ |
| Tubule Formation (<10 % vs ≥ 10%) | 1079 | 7.159 | $1.49 \times 10^{-12}$ |
| Nuclear Pleomorphism (3 vs 2 & 1) | 1079 | 20.777 | $4.40 \times 10^{-81}$ |
| Mitotic Count (>10 vs ≤10) | 1074 | 16.219 | $4.49 \times 10^{-53}$ |

**Supplementary Table 8. Hyperparameters Selected for Model Training.** Hyperparameters for recurrence score prediction are the result of Bayesian optimization for highest average tile area under the receiver operating characteristic curve over 50 iterations. Default hyperparameters established from training in other datasets were used for tumor identification.

| Hyperparameter | Recurrence Score Prediction | Tumor Identification |
|---|---|---|
| Augmentations | Flip, Rotate | Flip, Rotate, JPEG compression, Blur |
| Batch Size | 55 | 128 |
| Dropout | 0.31155932 | 0.5 |
| Epochs | 1 | 1 |
| Hidden Layer Width | 267 | 256 |
| Hidden Layers | 1 | 3 |
| L1 Loss | 0 | 0 |
| L1 Loss (Dense Layers) | 0.008306567 | 0 |
| L2 Loss | 0.034253246 | 0.00001 |
| L2 Loss (Dense Layers) | 0.039803477 | 0.00001 |
| Learning Rate | 0.000999282 | 0.0001 |
| Learning Rate Decay | 0.210307662 | 0.97 |
| Learning Rate Decay Steps | 1023 | 100000 |
| Loss | Mean Squared Error | Sparse Categorical Crossentropy |
| Model | Xception | Xception |
| Normalizer | Reinhard | Reinhard |
| Optimizer | Adam | Adam |
| Pooling | Average | Average |
| Tile Pixels | 299 | 299 |
| Tile um | 302 | 302 |
